# Sensitivity of Human Coronaviruses (HCoVs) to Heme Metabolism: Mechanisms Underlying Antiviral Effects

**DOI:** 10.64898/2026.09.30.755704

**Authors:** Noah T. Conohan, Kensuke Hirasawa

## Abstract

Heme metabolism regulates the availability of heme, an iron-containing molecule essential for oxygen transport, mitochondrial respiration, electron transfer, and enzymatic reactions. Increasing evidence implicates heme metabolism in regulating viral infection, potentially representing an important innate antiviral mechanism. Components of both heme biosynthesis and degradation pathways exhibit antiviral activity against different viruses; however, their contributions and conservation across human coronaviruses remain incompletely defined. We investigated the impact of modulating heme metabolism on infection by seasonal human coronaviruses HCoV-OC43 and HCoV-229E. Treatment with 5-aminolevulinic acid (5-ALA) produced cell line-dependent antiviral effects, with robust inhibition in H1299 cells and selective HCoV sensitivity in MRC-5 cells. These differences correlated with intracellular accumulation of protoporphyrin IX (PPIX), suggesting that cellular metabolic capacity influences antiviral efficacy. In contrast, components of the heme degradation pathway, including heme oxygenase-1 (HO-1), carbon monoxide, and biliverdin, did not exhibit antiviral effects against HCoV-OC43 or HCoV-229E. PPIX and synthetic porphyrin CoPPIX inhibited early infection through direct viral inactivation. Collectively, these findings indicate that antiviral effects of heme metabolism against seasonal human coronaviruses are primarily driven by direct inactivation by PPIX rather than cellular antiviral responses induced by the heme degradation pathway.

## Introduction

Human coronaviruses (HCoVs) are enveloped, positive-sense single-stranded RNA viruses that infect the human respiratory tract and cause disease ranging from mild illness to severe pneumonia. Seven HCoVs have been identified, including the alphacoronaviruses HCoV-229E and HCoV-NL63, and the betacoronaviruses HCoV-OC43, HCoV-HKU1, MERS-CoV, SARS-CoV, and SARS-CoV-2 [1]. The alphacoronaviruses 229E and NL63, along with the betacoronaviruses OC43 and HKU1, are seasonal coronaviruses that typically cause mild, cold-like symptoms. In contrast, MERS-CoV, SARS-CoV, and SARS-CoV-2 can cause severe disease with significant mortality. Although MERS-CoV (∼34%) and SARS-CoV (∼9.5%) exhibit higher case fatality rates, these viruses did not achieve the rapid and widespread global transmission observed during the COVID-19 pandemic caused by SARS-CoV-2 [2]. While vaccines against SARS-CoV-2 were authorized for emergency use in late 2020, 79.2 million cases and 1.7 million deaths had already been reported worldwide by the end of that year, highlighting the substantial delay between viral emergence and vaccine deployment [3]. These challenges underscore the need for understanding of host antiviral cellular pathways that may influence HCoV replication.

Heme, an essential molecule consisting of a porphyrin ring with a coordinated ferrous (Fe^2+^) ion at its center, is incorporated into many proteins as a prosthetic group to facilitate redox reactions. Hemoproteins participate in a wide range of biological processes, including gas transport, storage, and sensing, as well as catalysis and electron transport in metabolism [4]. In mammals, heme biosynthesis begins with the rate-limiting generation of 5-aminolevulinic acid (5-ALA), a non-proteinogenic amino acid, by aminolevulinic acid synthase (ALAS) [5] (Figure 1). Following a series of enzymatic reactions across the cytosolic and mitochondrial compartments, 5-ALA is converted to protoporphyrin IX (PPIX), the immediate precursor of heme. Insertion of a ferrous ion into PPIX by ferrochelatase (FECH) yields labile heme, which is subsequently incorporated into hemoproteins or degraded by constitutive heme oxygenase-2 (HO-2) to mitigate cytotoxicity. During oxidative stress or increased levels of intracellular labile heme, an inducible isoform of heme oxygenase (HO-1) is expressed [6]. The degradation of heme results in the production of carbon monoxide, ferrous iron, and biliverdin. Biliverdin is reduced to bilirubin by biliverdin reductase (BVR), which is then exported from cells.

**Figure 1.**
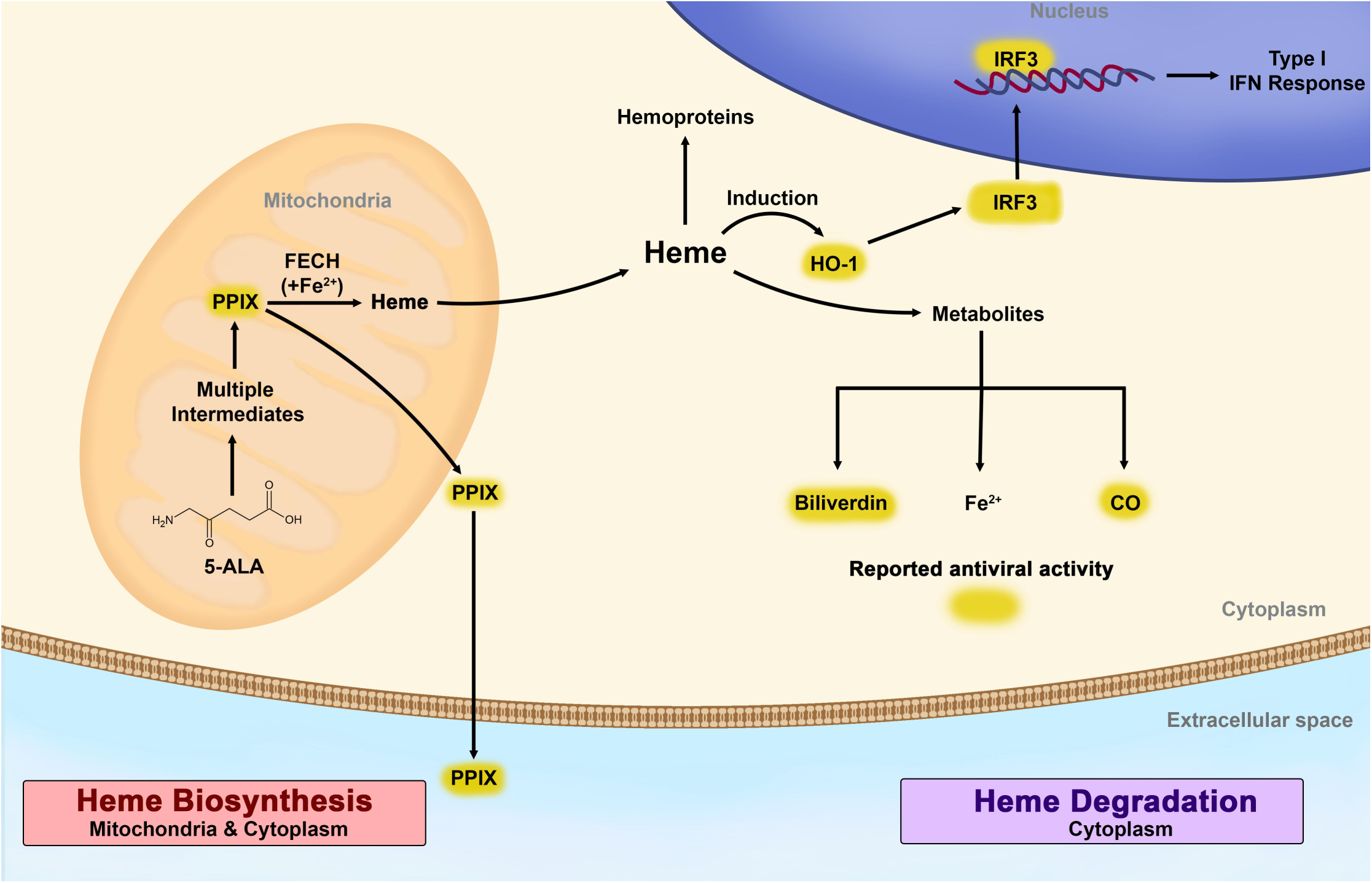
Overview of heme metabolism pathways and their roles in innate antiviral responses. Schematic representation of heme metabolism and its potential roles in innate antiviral responses. The heme biosynthetic pathway proceeds from 5-aminolevulinic acid (5-ALA) through a series of intermediates to generate protoporphyrin IX (PPIX), which is converted to heme by incorporation of iron. Heme induces heme oxygenase-1 (HO-1), which promotes heme degradation into biliverdin (BV), carbon monoxide (CO), and free iron. Heme can also be incorporated into heme-containing proteins. Pharmacologic modulators such as CoPPIX can be used to induce expression of HO-1. PPIX, HO-1-mediated activation of IRF3, BV, and CO (highlighted in yellow) have been reported to exert antiviral activities, suggesting that multiple components of the heme metabolic pathway contribute to innate antiviral responses.

Beyond its role in cellular metabolism, heme homeostasis has been increasingly recognized as a modulator of antiviral responses. Along the biosynthetic axis, exogenous administration of 5-aminolevulinic acid (5-ALA) promotes intracellular accumulation of protoporphyrin IX (PPIX), which has demonstrated antiviral activity in multiple replication systems, including coronaviruses and pestiviruses [7–9]. Consistently, direct administration of PPIX inhibits infection across a range of enveloped viruses, including vesicular stomatitis virus, through mechanisms involving disruption of viral entry and replication [10–12]. Downstream, heme oxygenase-1 (HO-1) has been the subject of investigation as a modulator of innate immunity and cellular stress that may influence viral replication [13–18]. The antiviral effects of heme oxygenase-1 (HO-1) has been attributed to both the activity of its metabolites and its ability to modulate interferon signaling pathways, depending on the viral context. In studies investigating influenza A virus (IAV), respiratory syncytial virus (RSV), and spring viremia of carp virus (SVCV), HO-1 is involved in enhancing interferon responses, including activation of interferon regulatory factor 3 (IRF3) [16–18]. In addition to HO-1, metabolites biliverdin and carbon monoxide appear to have antiviral effects through modulation of redox status, direct binding, or indirect signalling [13–15]. Taken together, both axes of heme homeostasis represent viable candidates to for host-directed antiviral strategies.

Despite evidence that modulation of heme metabolism can influence coronavirus infection, these effects have primarily been characterized in SARS-CoV-2 and feline infectious peritonitis virus (FIPV) [7,8] and it remains unclear whether they are conserved across human coronaviruses. Moreover, the specific components of heme metabolism responsible for mediating antiviral responses against human coronaviruses (HCoVs) have not been fully defined. Here, we investigate how elements of heme metabolism can inhibit HCoVs using seasonal coronaviruses, HCoV-229E and HCoV-OC43.

## Methods

### Cells, viruses, and reagents

The human lung fibroblast cell line MRC-5 and human epithelial lung cancer cell line H1299 were purchased from the American Type Culture Collection (ATCC, Manassas, VA, USA). Human coronaviruses OC43 (HCoV-OC43) and 229E (HCoV-229E) were obtained from the American Type Culture Collection (ATCC). 5-aminolevulinic acid hydrochloride was purchased from Sigma-Aldritch (Milwaukee, WI, USA). Concentrated stock solutions were prepared by dissolving 5-ALA in culture media and stored at −80℃ except for PPIX quantification assays, in which freshly reconstituted 5-ALA was used. Cobalt protoporphyrin (III) chloride (CoPPIX) and biliverdin hydrochloride (BV) were obtained from Frontier Specialty Chemicals (Logan, UT, USA). Stock solutions were freshly prepared by dissolving compounds in DMSO and then diluted to working concentrations in culture media. CORM3 was purchased from Tocris Bioscience (Bristol, UK) and reconstituted in distilled water and diluted in culture media immediately prior to use. Inactivated CORM3 (iCORM3) was generated by reconstituting CORM3 in PBS and incubating for 24 hours at room temperature. Scrambled/Negative control siRNA and siRNA targeting HO-1 (s194530) were purchased from Life Technologies (Carlsbad, CA, USA).

### Cell culture

All cells were cultured in high-glucose Dulbecco’s modified Eagle’s medium (Corning, MA, USA) supplemented with 10% fetal bovine serum (HyClone, Cytiva, USA), 1 mM sodium pyruvate (Life Technologies, Carlsbad, CA, USA) and antibiotic-antimycotic (Thermo Scientific, Waltham, MA, USA). Cells were maintained in 10-cm culture dishes at 37°C with 5% CO_2_ for the use of experiments in this work.

### Viral infection

Cells were plated were plated in 24-well plates (8×10^4^ cells/well) and left overnight to attach. The following day, cells were infected with human coronavirus 229E or OC43 at an MOI of 0.01. The diluted stock viruses were adsorbed for 2 hours at 33°C in serum-free DMEM, which was then removed and replaced with infection media (DMEM with 2% FBS). Infected cells were incubated at 33°C with 5% CO_2_. For siRNA knockdown, cells were transfected with 5 pmol siRNA oligo using Lipofectamine RNAiMAX Transfection Reagent (Life Technologies, Carlsbad, CA, USA) added to culture media 24 hours prior to infection. Treatment with 5-ALA was administered in culture media 4 hours prior to infection to induce PPIX accumulation prior to infection, whereas other compounds were added in infection media following adsorption (2 HPI) to assess their ability to impact replication.

### Cell viability assay

Viability of cells treated with 5-ALA was measured using the Colorimetric Cell Viability Kit I (WST-8) following manufacturer’s instructions (PromoCell GmbH, Germany). After 24 hours of treatment with 5-ALA or CORM3, 10 µl of reagent was added to each well of 96-well plates containing cells (100 μl media) and incubated in the dark at 37°C for 40 minutes. Absorbance was measured at 450 nm using a Bio-Rad Model 3550 microplate reader and cell viability was determined based on comparison to untreated controls. For CoPPIX and BV, cell viability was determined by crystal violet assay. Following 24 hours of treatment with CoPPIX or biliverdin, cells were washed with PBS twice and subsequently fixed with 10% buffered formalin solution for 1 hour, followed by a second wash step and 1 hour stain with 0.5% crystal violet solution. Fixed and stained plates were then washed and decanted, before allowing to dry overnight. The following day, 100 μl of 30% acetic acid was added to each well and the plate was shaken for 5 minutes prior to measurement. Absorbance was measured at 595 nm using an Agilent BioTek Epoch microplate reader (Santa Clara, CA, USA).

### Western blot analysis

Samples for western blot analysis of proteins were collected and processed as previously described [19]. In this study, the following antibodies were used: α-229E nucleocapsid protein from Ingenasa (Madrid, Spain), α-OC43 nucleocapsid protein (MAB9013) from MilliporeSigma, α-HO-1 (ab68477) from AbCam (Cambridge, UK), and GAPDH (sc-365062) from Santa Cruz Biotechnology (Dallas, USA). Secondary antibodies α-mouse-HRP (#1706516) and α-rabbit-HRP (#1706515) were purchased from Bio-Rad Laboratories (Hercules, CA, USA). Membranes were exposed and images captured using a Bio-Rad ChemiDoc^TM^ imaging system (Hercules, CA, USA).

### PPIX quantification

MRC-5 and H1299 cells were plated separately in 24-well plates (6 x 10^4^ cells/well) and incubated overnight to attach. Following this, cells were treated with different concentrations (0.625, 1.25, and 2.5 mM) of 5-ALA and incubated at 37℃ in the dark for 4 hours. After incubation, culture supernatants and lysate samples were collected and stored immediately at −80℃ in opaque amber tubes. Lysate samples were obtained by washing cells with PBS before lysis with supplemented RIPA buffer (RIPA, PMSF, aprotinin, and Halt^TM^ phosphatase and protease inhibitor cocktail (Thermo Scientific)). In minimally lit conditions, lysate samples were diluted in 1:40 in PBS supplemented with Triton X-100 to avoid aggregation of PPIX (Triton X-100 0.1% in PBS). Likewise, supernatant was diluted 1:10 using Triton X-100 0.1% in PBS. Diluted samples were then added to solid black 96-well plates (Fluotrac, Greiner Bio-one), covered with foil, and protected from light until measurement. Fluorescence of PPIX was measured with a Biotek Synergy MX plate reader with an excitation wavelength of 405 nm and emission wavelength of 635 nm. Wells with appropriately diluted unconditioned media and lysis buffer were used as blank controls.

### Viral neutralization/inactivation assays

11 200 PFU of OC43 or 19 600 PFU of 229E was incubated with 50 μl of CoPPIX at indicated concentrations at for 1h at 33℃ in the dark. Treated virus was then serially diluted in serum-free culture media and adsorbed for 2h. Following adsorption, cells were washed twice with PBS and media was replaced with 2% FBS DMEM. Plates were incubated for 7 days, and infection was measured as described for TCID_50_ (50% tissue culture infective dose) assay [20]. In instances where one or more replicates were below the limit of detection, they were labelled “Not detected” (ND) or assigned a value of 0 if other replicates could be quantified.

### Statistical analyses

Statistical analyses were performed using Prism 7.0 (GraphPad). Two-way ANOVA with Šídák’s posthoc test was used to compare multiple groups. Dunnett’s posthoc test was used for comparisons to a single group. *P* < 0.01 were considered statistically significant.

### Data availability

The datasets used and/or analysed during the current study available from the corresponding author on reasonable request.

## Results

To begin, we wanted to determine the sensitivity of HCoV OC43 and 229E to antiviral activities of the heme biosynthesis axis. We determined working concentrations of 5-ALA in MRC-5 (normal human lung fibroblasts) and H1299 (human non-small cell lung carcinoma) by cell viability assay (Figure 2A). Next, we treated cells with different concentrations of 5-ALA for 4 hours prior to OC43 or 229E infection to permit accumulation of PPIX or activate downstream elements. Cell lysates were collected after 1- and 2-days post infection and infection was assessed by western blot analysis. We observed that 5-ALA pretreatment had no effect on viral replication of OC43 in MRC-5 cells at 1- and 2-days post infection (DPI) (Figure 2B). In contrast, OC43 replication was impaired at 1 DPI in H1299 cells treated with 5-ALA as demonstrated by decreases in viral nucleocapsid expression. When the same conditions were applied to 229E infection, 5-ALA suppressed replication of 229E at 2 DPI in MRC-5 cells (Figure 2C). Moreover, in H1299 cells, N protein expression was observed in infected cells without treatment, but not in those treated with 5-ALA (0.025 to 10 mM) at 1 DPI. A marked dose-dependent decrease with 5-ALA that was also observed at 2 DPI. These results indicate that antiviral effects of 5-ALA are context dependent based on both cell line and virus, as 229E showed sensitivity in both cells line, but OC43 infection was only sensitive to 5-ALA treatment in H1299 cells, but not MRC-5.

**Figure 2.**
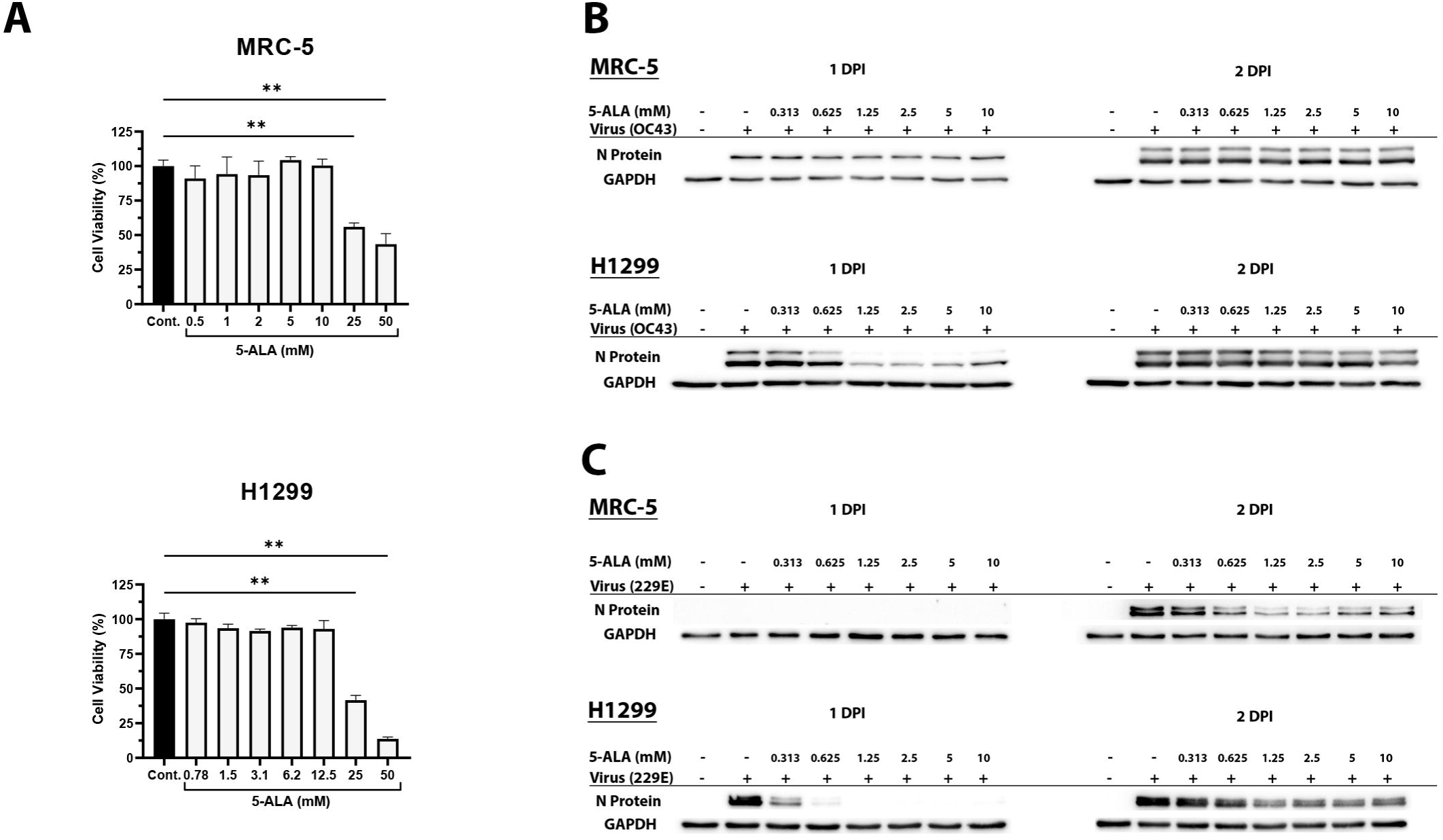
Administration of exogenous 5-ALA leads to reduced viral infection. (A) The effect of 5-ALA on cell viability was determined in MRC-5 and H1299 cells using WST-8 assay. (B–C) MRC-5 (B) and H1299 (C) cells were treated with 5-ALA at the indicated concentrations and times beginning 4 h prior to infection with HCoV-229E or HCoV-OC43 at an MOI of 0.01. Viral infection was assessed at 1- or 2-days post-infection by western blot analysis using antibodies against OC43 or 229E nucleocapsid protein (N) and GAPDH. Western blots cropped for display purposes.

To investigate the differences in antiviral effects of 5-ALA in our system, we sought to examine the ability of each cell line to metabolize 5-ALA into protoporphyrin (PPIX), which has been recognized to have antiviral capacity [10–12]. We treated an equal number of MRC-5 and H1299 cells with 5-ALA for 4 hours and subsequently collected supernatant and lysate samples to assess extracellular and intracellular PPIX content by fluorometric measurement (Figure 3). H1299 produced higher levels of both extracellular and intracellular PPIX than MRC-5, which indicated that accumulation of antiviral PPIX may explain differences in antiviral capabilities of 5-ALA between both cell lines.

**Figure 3.**
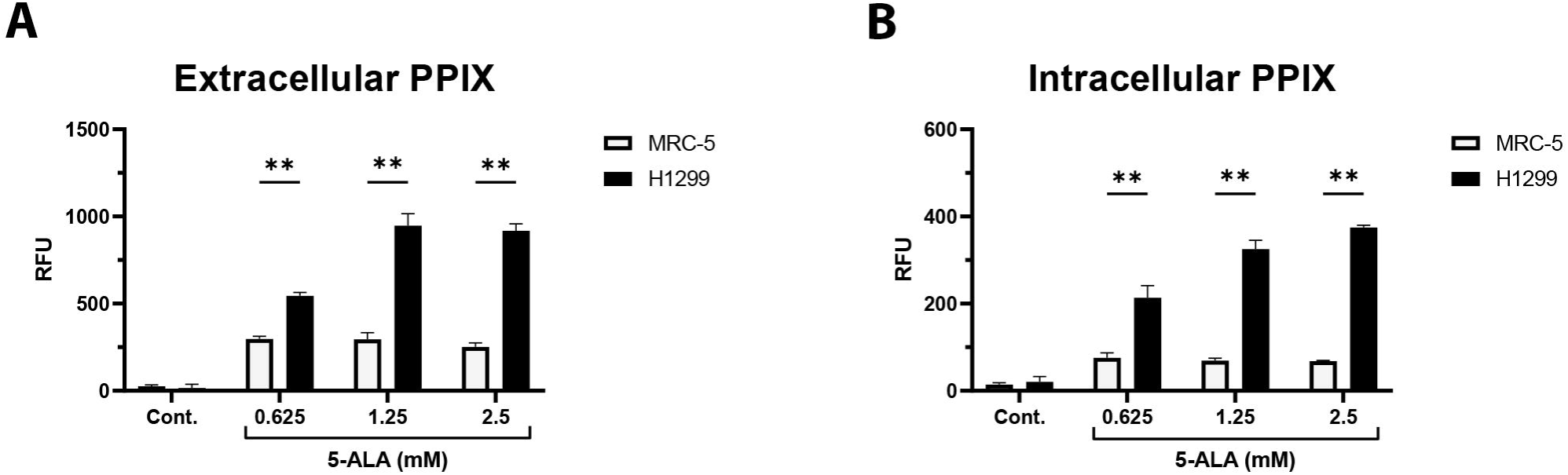
5-ALA treatment induces greater PPIX production and accumulation in H1299 cells than in MRC-5 cells. Intracellular and extracellular protoporphyrin IX (PPIX) levels were measured following 4 hours of incubation with 5-ALA by fluorimetry of lysate and supernatant samples respectively (Ex/Em: 405/635 nm). Data are presented as [mean ± SEM] of Relative Fluorescence Units (RFU) **p<0.01 by two-way ANOVA with Šídák’s post-hoc test.

We next examined the sensitivity of HCoV OC43 and 229E to the degradation axis of heme metabolism. To do this, we utilized the synthetic metalloporphyrin cobalt (III) protoporphyrin IX (CoPPIX) as a potent inducer of the cytoprotective enzyme heme oxygenase-1 (HO-1). HO-1 catalyzes the oxidative cleavage of heme to generate CO, Biliverdin and Fe^2+^ (Figure 1). We used CoPPIX concentrations up to 25 µM, as higher doses induced cytotoxicity (Figure 3A). When MRC-5 cells were treated with CoPPIX at 2 hours before OC43 infection, HO-1 expression was induced, and infection was inhibited at a concentration of 25 µM (Figure 3B). Next, we sought to functionally evaluate if the induction of HO-1 contributes to the inhibition of HCoV infection by CoPPIX. MRC-5 cells were treated with control (scrambled) or HO-1 siRNA for 24 hours prior to infection and subsequent treatment with or without CoPPIX (Figure 3C). In both OC43 and 229E infection, OC43 or 229E replication was inhibited in MRC-5 cells treated with CoPPIX. Moreover, HO-1 siRNA effectively reduced CoPPIX-induced HO-1 expression. However, HO-1 knockdown did not alter the antiviral effect of CoPPIX, as viral protein remained below detection in all treated samples regardless of HO-1 expression. These results indicate that HO-1 and its downstream elements are not involved in antiviral activity of heme metabolism against HCoVs. Biliverdin (BV), a downstream metabolite of heme degradation, has documented antiviral capacity in other viral contexts (13-15). To determine the sensitivity of HCoVs OC43 and 229E to BV, we first conducted cell viability assays to access the cytotoxic effects of biliverdin in MRC-5 (Figure 5A). When we tested antiviral activity of BV to HCoVs, we found that treatment with exogenous biliverdin did not impact viral replication of HCoV OC43 or 229E in MRC-5 cells (Figure 5B). Lastly, to determine antiviral activity of another downstream product of heme degradation, carbon monoxide (CO), we used CORM-3, a carbon monoxide–releasing molecule (21). We first defined a working concentration of CORM-3 (Figure 6A). Treatment of infected cells with CORM3 impaired production of HCoV 229E viral protein, but this effect cannot be attributed to CO release because the effect was also observed in inactivated CORM3 (iCORM3), which has been depleted of CO (Figure 6B). Taken together, these results suggest that HCoVs OC43 and 229E are not sensitive to the antiviral components of the heme degradation pathway.

Several porphyrin molecules have been reported to interact directly with viral particles and reduce infectivity (10, 22). To determine whether this is also the case for HCoVs, we performed neutralization assays with porphyrin molecules using both HCoV-OC43 and −229E (Figure 7). The viruses were incubated with different concentrations of PPIX (25-200 mM), CoPPIX (25-200 mM), or vehicle control (DMSO) in the absence of cells at 33℃ for 1 hour. Then viruses were serially diluted, adsorbed with MRC-5 cells for 2 hours, and washed with PBS prior to media replacement. As shown in Figure 7, infectious viral particles of HCoV-OC43 and HCoV-229E were significantly reduced when preincubated with PPIX at a concentration of 50 μM (Figure 7A). Moreover, no infectious viral particles of HCoV-OC43 or HCoV-229E were detected at PPIX concentrations of 100 and 200 μM. Similarly, CoPPIX pretreatment reduced infectivity of HCoV-OC43 and HCoV-229E starting at concentrations of 100 μM and 50 μM, respectively (Figure 7B). These results indicate that antiviral effects of heme metabolism against HCoVs OC43 and 229E are mediated by porphyrins such as PPIX, rather than cell-mediated interactions through HO-1 induction or downstream metabolites biliverdin or CO.

## Discussion

Heme metabolism has emerged as a potential modulator of viral infection. Studies suggest antiviral roles for elements of heme biosynthesis and degradation across multiple virus families, notably enveloped RNA viruses of the Flaviviridae and Coronaviridae families [7–17]. However, antiviral mechanisms induced by heme metabolism, and the extent to which they are conserved across human coronaviruses, remain incompletely defined. In this study, we demonstrate that the seasonal coronaviruses HCoV-OC43 and HCoV-229E are sensitive to the antiviral effects of heme biosynthesis, which are caused by direct disruption of the viral particle by protoporphyrin IX (PPIX), an intermediate of heme biosynthesis.

Our results indicate that 229E is sensitive to 5-ALA treatment in both MRC-5 and H1299 cells, while OC43 only exhibits sensitivity during infection of H1299 (Figure 2). Moreover, we demonstrated that H1299 treated with 5-ALA featured increased intracellular accumulation and export of PPIX compared to MRC-5 (Figure 3). Transformation of cells enhances heme biosynthesis activity, leading to increased PPIX production upon stimulation with 5-ALA (23). This may explain the discrepancy between cell lines, as H1299 cells are human non-small cell lung carcinoma cells, whereas MRC-5 cells are normal human lung fibroblasts. These findings suggest that variability in heme biosynthetic capacity among cell lines may represent an important, and often overlooked, confounding factor in antiviral studies of heme biosynthesis. Notably, many *in vitro* replication systems rely on cancer-derived cell lines, in which heme biosynthesis is frequently dysregulated, potentially leading to enhanced PPIX accumulation and exaggerated antiviral responses [24]. In contrast, primary or non-transformed respiratory cell lines, such as MRC-5 cells, may more accurately reflect physiological conditions but exhibit reduced capacity for PPIX accumulation. As such, differences in heme metabolism may influence both the magnitude and reproducibility of antiviral effects mediated by 5-ALA treatment. Despite this limitation, our findings also suggest that 5-ALA supplementation may be effective in cellular contexts with high capacity for PPIX production, such as some myeloid-derived cells [25,26]. Consistent with this, antiviral effects of 5-ALA have been reported in viruses with tropism for myeloid-derived cells, including feline infectious peritonitis virus (FIPV) and classical swine fever virus (CSFV), with *in vivo* studies against the latter demonstrating protective effects following 5-ALA administration [7,9].

Extracellular virucidal activity of PPIX and related porphyrins has been described in several viral systems. This is through the ability of these compounds to disrupt viral envelope, preventing viral entry or interactions with host cell membranes [10–12,23,27]. However, while direct extracellular inactivation by PPIX is a well-established event, the mechanisms underlying intracellular inhibition of viral replication remain less well defined. In a 2024 study, Doki et al. demonstrated that 5-ALA co-administered with iron donor SFC promoted conversion of intracellular PPIX to heme and partially rescued viral replication of alphacoronavirus FIPV, suggesting intracellular PPIX may directly impact viral replication [7]. One potential explanation involves the ability of protoporphyrin IX to interact with G-quadruplex (G4) structures in RNA [28–30]. Given that coronavirus genomes contain G4-forming sequences that vary between species, differential interactions with PPIX may contribute to virus-specific sensitivity, although this has yet to be explicitly investigated [31,32]. In this study, we found that PPIX and CoPPIX directly inactivate the infectivity of HCoVs (Figure 7), suggesting that porphyrins can impact HCoV replication through extracellular effects on viral particles. In addition, both intra- and extracellular induction of PPIX by 5-ALA effectively inhibited HCoV infection (Figure 2 and 4), which may reflect the combined antiviral effects of PPIX within and outside infected cells. Collectively, these findings highlight the importance of PPIX in antiviral responses against HCoVs, although further investigation into the intracellular mechanisms by which PPIX disrupts viral replication is warranted.

**Figure 4.**
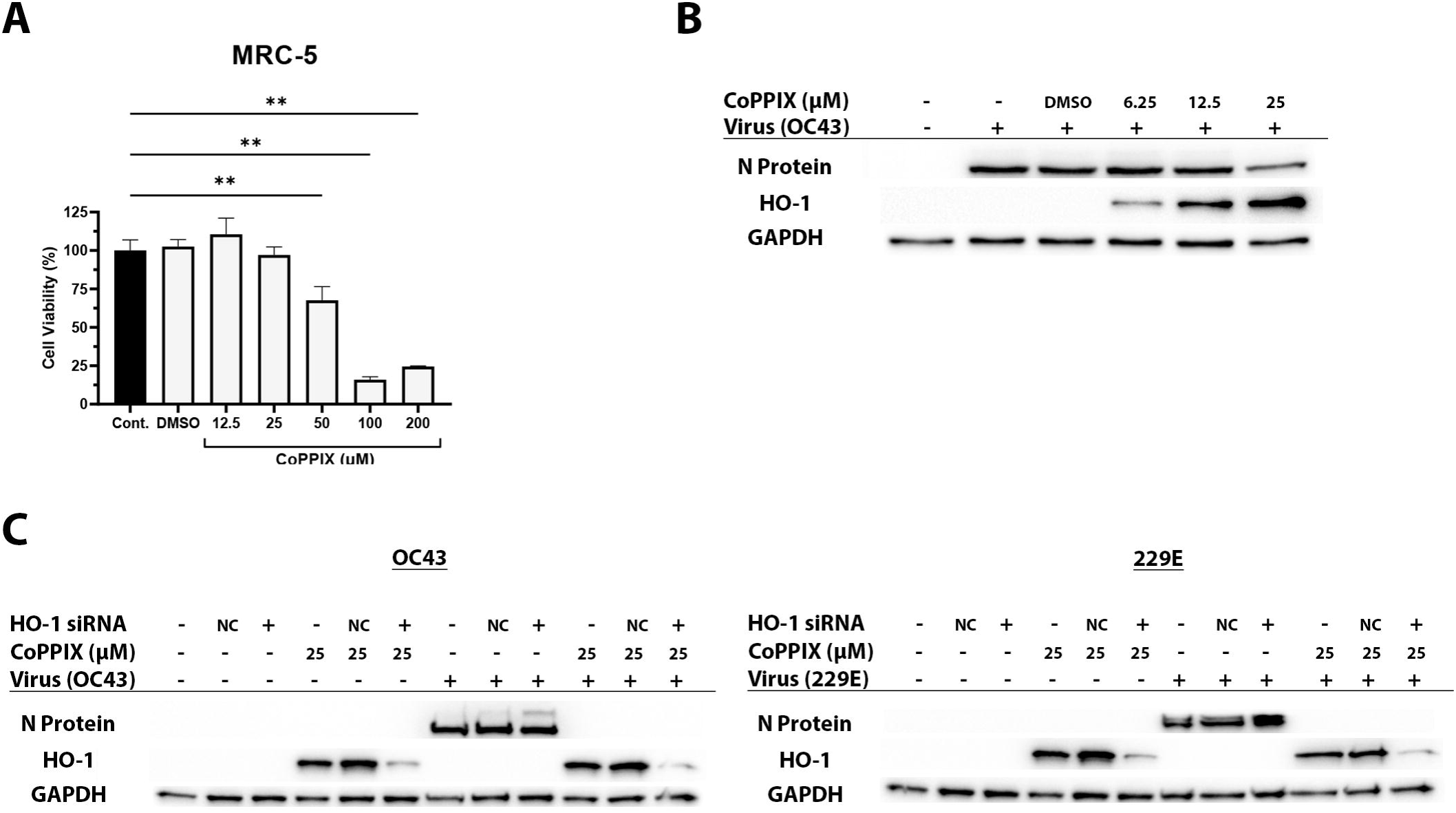
Cobalt Protoporphyrin IX inhibits human coronavirus infection independently of heme oxygenase-1. (A) The cytotoxicity of cobalt protoporphyrin IX (CoPPIX) in MRC-5 cells was determined via crystal violet assay. (B) MRC-5 cells were treated with at the indicated concentrations of CoPPIX 2 hours post-infection with HCoV-OC43 or HCoV-229E. (C) To assess involvement of HO-1, siRNA knockdown was completed 24 hours prior to infection and subsequent CoPPIX treatment. Viral infection was assessed at 1-day post-infection by western blot analysis using antibodies against HO-1, OC43 N protein, 229E N protein or GAPDH. **p<0.01 by one-way ANOVA with Dunnett’s post-hoc test. Western blots cropped for display purposes.

In addition to heme biosynthesis, we also explored the sensitivity of OC43 and 229E to the heme degradation pathway, as it has been widely investigated for its potential in shaping antiviral responses [33]. Primarily recognized for its cytoprotective role during oxidative stress, HO-1 has also been shown to play a role in interferon responses during viral infection. Specifically, independent of its catalytic activity, HO-1 has been shown to promote IRF3 phosphorylation and nuclear translation during influenza A virus infection of RAW264.7 cells [16]. Furthermore, a functional study by Wu et al. revealed that HO-1 plays bidirectional role, promoting type I interferon signalling through IRF3/7 stabilization during spring viremia of carp virus (SVCV) infection, while promoting degradation during oxidative stress [18]. Notably, we previously reported that HCoV-OC43 and 229E are sensitive to IRF3-mediated antiviral responses (19). Nevertheless, in this study, HO-1 knockdown did not reduce antiviral activity of the heme biosynthesis (Figure 4), suggesting that activation of IRF3 by HO-1 may not play a significant role in antiviral responses during seasonal HCoV infection. We did not also find antiviral activities of BV and CO against the HCoVs (Figure 5 and 6). Interestingly, viruses such as Dengue virus (DENV) and SARS-CoV-2 have been shown to inhibit of regulators of antioxidant signalling upstream of HO-1, such as Nrf2 and Sirt1, suggesting that oxidative stress favors replication of these viruses [34,35]. To this end, biliverdin acts a scavenger of reactive oxygen species and has been known to bind to viral proteins [15,36]. In contrast to our results, biliverdin was shown to greatly suppress SARS-CoV-2 replication despite belonging to the same betacoronavirus genus as HCoV-OC43 [34]. This further emphasizes the diversity of HCoVs and how it can impact their susceptibility to antiviral components of heme metabolism.

**Figure 5.**
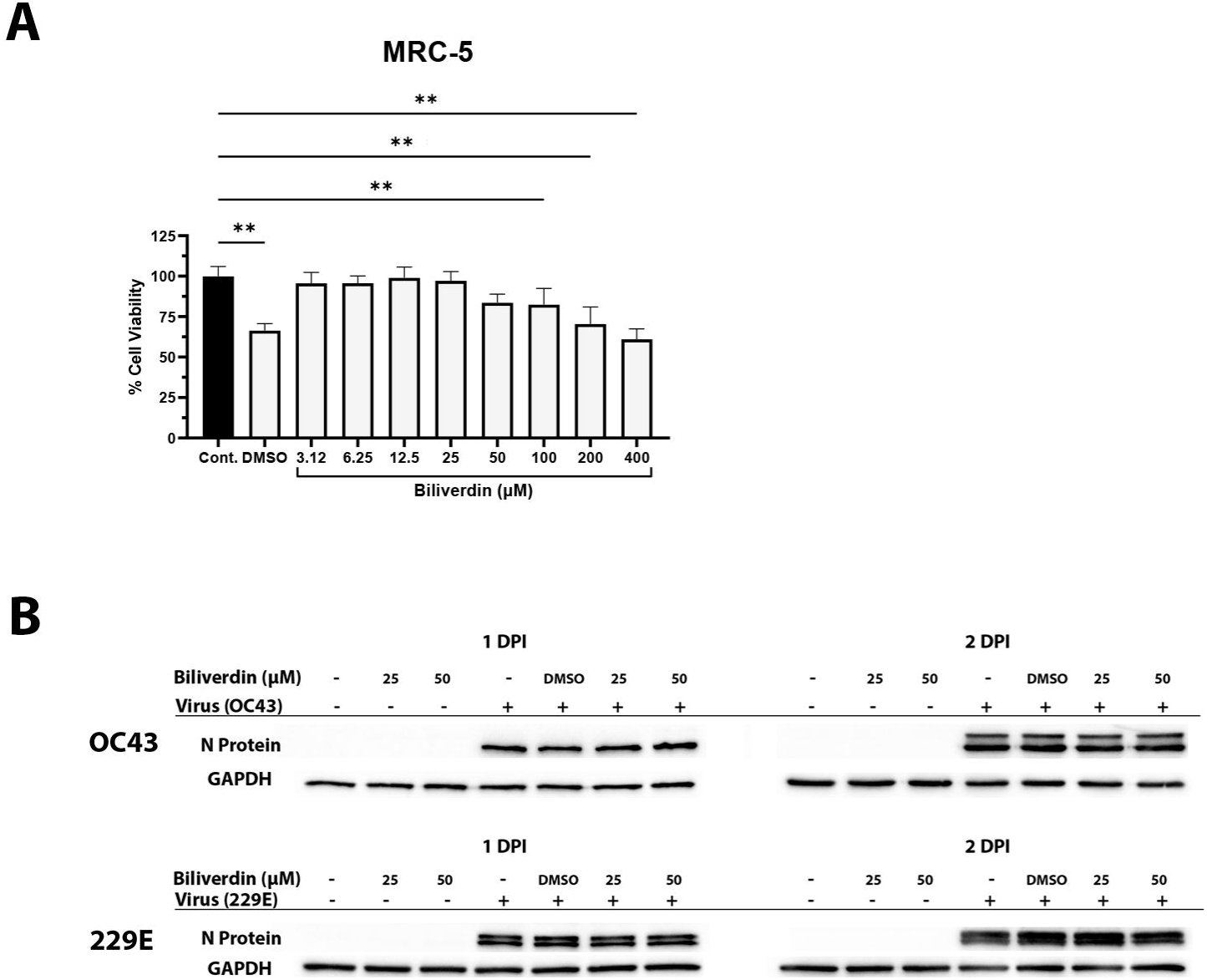
Heme oxygenase metabolite biliverdin does not inhibit HCoV-OC43 or HCoV-229E infection. (A) Cytotoxicity of biliverdin in MRC-5 cells was determined via crystal violet assay. DMSO vehicle control is matched to highest concentration tested (400 μM). (B) MRC-5 cells were treated with at the indicated concentrations of biliverdin 2 hours post-infection with HCoV-OC43 or HCoV-229E. Viral infection was assessed at 1- or 2-days post-infection by western blot analysis using antibodies against OC43 or 229E nucleocapsid protein (N) and GAPDH. **p<0.01 by one-way ANOVA with Dunnett’s post-hoc test. Western blots cropped for display purposes.

**Figure 6.**
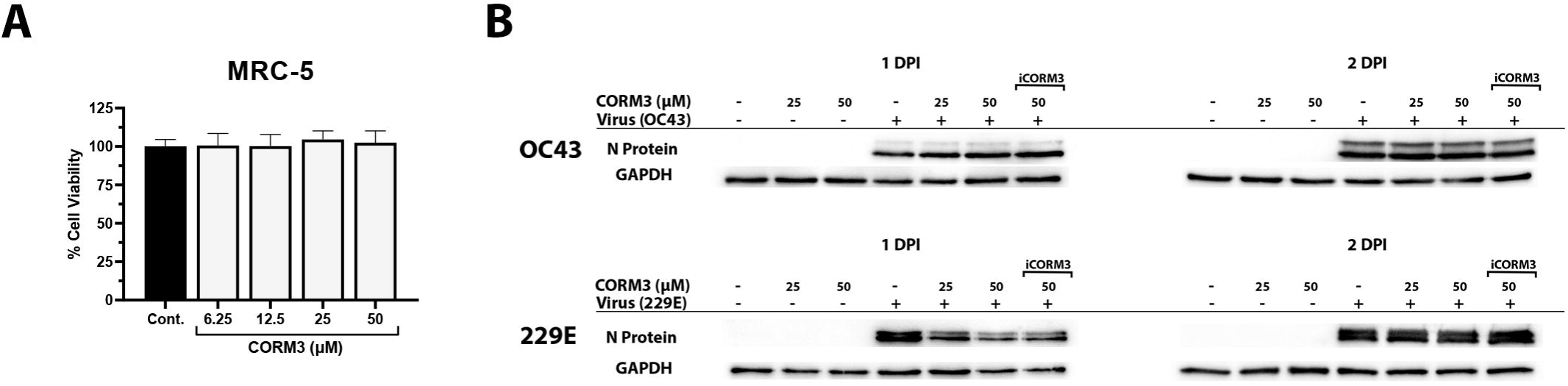
Carbon monoxide donor CORM3 does not inhibit HCoV infection. (A) The cytotoxicity of CORM3 in MRC-5 cells was determined via WST-8 assay. (B) MRC-5 cells were treated with at the indicated concentrations of CORM3 or inactivated CORM3 (iCORM3) 2 hours post-infection with HCoV-OC43 or HCoV-229E. Viral infection was assessed at 1- or 2-days post-infection by western blot analysis using antibodies against OC43 or 229E nucleocapsid protein (N) and GAPDH. **p<0.01 by one-way ANOVA with Dunnett’s post-hoc test. Western blots cropped for display purposes.

**Figure 7.**
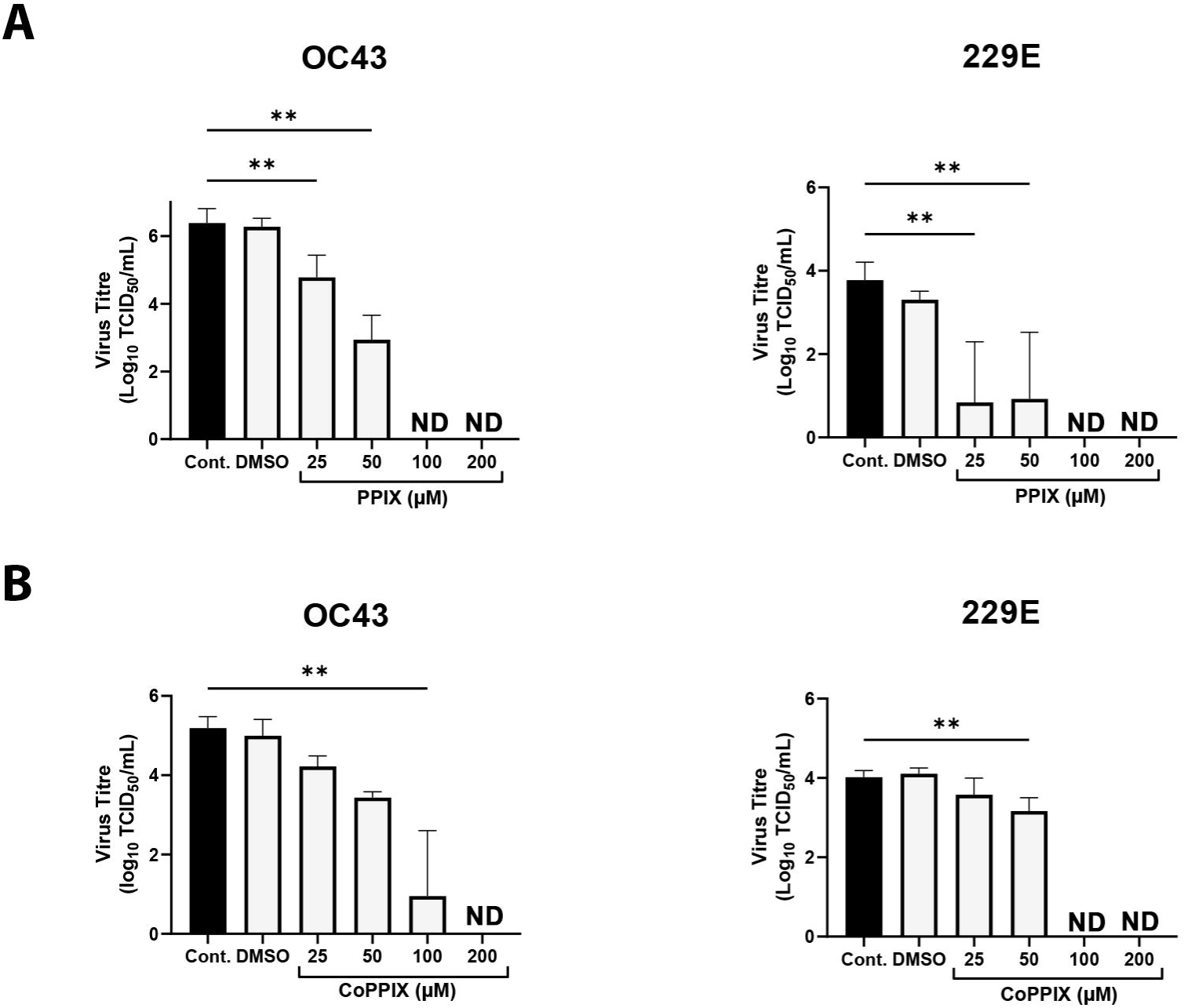
Porphyrins directly reduce viral infectivity. (A) Viral particles were incubated with protoporphyrin IX (PPIX) or (B) cobalt protoporphyrin IX (CoPPIX) at the indicated concentrations for 1h at 33°C prior to adsorption. Following adsorption, cells were washed, and media was replaced. At 7 days post-infection, cytopathic effect was recorded, and viral titres were determined by TCID50 assay. Titre values the below limit of detection are indicated as “ND”. **p<0.01 by one-way ANOVA with Dunnett’s post-hoc test.

In conclusion, our study is the first to demonstrate that heme biosynthesis possesses antiviral activity against seasonal HCoVs. We further show that this antiviral activity is mediated by the direct virucidal action of PPIX against HCoVs. These findings suggest that PPIX or its derivatives may represent promising virucidal agents for antiviral therapeutics and environmental disinfection of seasonal HCoVs.

## Acknowledgements

We thank Ariyana Hasan and Adam Salman for their technical assistance processing samples for western blot analysis.

This work was supported by Memorial University’s Medical Research Foundation Grant.

## Author Contributions Statement

Conceptualization/Design: NC, KH; Data Collection: NC; Data Analysis: NC, KH; Writing original draft: NC, KH. Review and Editing: NC, KH.

## Additional Information

The authors declare no competing interests.

